# A two-hit adversity model in developing rats reveals sex-specific impacts on prefrontal cortex structure and behavior

**DOI:** 10.1101/2020.10.23.352161

**Authors:** Kelsea R. Gildawie, Lilly M. Ryll, Jessica C. Hexter, Shayna Peterzell, Alissa A. Valentine, Heather C. Brenhouse

**Affiliations:** Department of Psychology, Northeastern University, Boston, MA, USA

**Keywords:** Maternal separation, prefrontal cortex, perineuronal nets, parvalbumin, sex differences, anxiety

## Abstract

Adversity early in life substantially impacts prefrontal cortex (PFC) development and vulnerability to later-life psychopathology. Importantly, repeated adverse experiences throughout childhood increase the risk for PFC-mediated behavioral deficits more commonly in women. Evidence from animal models points to effects of adversity on later-life neural and behavioral dysfunction; however, few studies have investigated the neurobiological underpinnings of sex-specific, long term consequences of multiple developmental stressors. We modeled early life adversity in rats via maternal separation (postnatal day (P)2-20) and juvenile social isolation (P21-35). Adult (P85) male and female rats were assessed for differences in the presence and structural integrity of PFC perineuronal nets (PNNs) enwrapping parvalbumin (PV)-expressing interneurons. PNNs are extracellular matrix structures formed during critical periods in postnatal development that play a key role in the plasticity of PV cells. Females – but not males – exposed to multiple hits of adversity demonstrated a reduction in PFC PV cells in adulthood. We also observed a sex-specific, potentiated reduction in PV+ PNN structural integrity. Moreover, correlations between neural disruption and hyperactivity/risk-assessment behavior were altered by adversity differently in males and females. These findings suggest a sex-specific impact of repeated adversity on neurostructural development and implicate PNNs as a contributor to associated behavioral dysfunction.

## 1. Introduction

Adverse environmental influences during sensitive periods of development can disrupt prefrontal cortex (PFC) maturation (Chocyk et al., 2010; Stevenson et al., 2008) and increase vulnerability to later-life onset of neuropsychiatric disorders, including schizophrenia, post-traumatic stress disorder, and anxiety disorders (Agid et al., 1999; Heim et al., 2008, 2009). Importantly, evidence suggests that adversity early in life increases the likelihood of further adverse experiences throughout development (Manyema et al., 2018). Clinical studies demonstrate enhanced vulnerability to a second “hit” of adversity (McLaughlin et al., 2010), where repeated stressful life events throughout childhood result in more severe neuropsychiatric symptoms, particularly in women (Bale & Epperson, 2015; Manyema et al., 2018; Suliman et al., 2009). There is, therefore, a critical need to understand the sex-specific neuromolecular underpinnings of developmental responses to adversity, and how this may relate to dysfunction later in life.

Several paradigms exist to model early life adversity in rodents (Molet et al., 2014; Schmidt et al., 2011), improving our understanding of neural mediators driving the onset of behavioral disruption (Sarkar et al., 2019). The maternal separation (MS) paradigm is a well-characterized analogue to childhood neglect in humans (Lehmann & Feldon, 2000). Repeated isolation from the dam throughout postnatal development results in increased anxiety-like behavior (Daniels et al., 2004; Ganguly et al., 2015), which has been seen to affect females more than males in adolescence (Honeycutt et al., 2020). Isolation paradigms have also been utilized after postnatal development to elicit behavioral disruption later in life (see review: Fone & Porkess, 2008). While social isolation (SI) has often been employed to induce adversity throughout life after weaning (Lapiz et al., 2003; Liu et al., 2019; Möller et al., 2013a; Schiavone et al., 2009), few studies report SI during discrete developmental time points, such as juvenility (Bicks et al., 2020; Pietropaolo et al., 2008) or different periods of adolescence (Jaric et al., 2019; Li et al., 2018). Evidence regarding sex-dependent effects of SI is also sparse, with inconsistent effects on anxiety-like behavior in males and females (Pietropaolo et al., 2008; Rodgers & Cole, 1993). Notably, early life adversity paradigms are often combined to assess the additive effects of multiple adverse experiences throughout development (Castillo-Gómez et al., 2017; Deslauriers et al., 2013; Jaric et al., 2019; Monte et al., 2017; Rincel et al., 2019). Some evidence suggests that there are sex-specific effects of multiple hits of developmental stressors on later-life anxiety-like behavior, working memory, and social behavior (Hudson et al., 2014; Monte et al., 2017; Rincel et al., 2019); however, a dearth of information remains regarding the potential neuroplastic mediators underlying sex-specific effects of multiple behavioral stressors on anxiety-like behavior.

The neonatal (Greenhill et al., 2015) and juvenile (Bicks et al., 2020) periods are implicated as stages of increased plasticity and susceptibility to disrupted environment. These windows of susceptibility have been linked to the maturation of cortical excitatory/inhibitory balance (Hensch, 2004, 2005). GABAergic machinery maturation is dynamic throughout early life (Minelli et al., 2003); however, the structural development of GABAergic circuitry occurs chiefly during early life, concluding towards the end of juvenility (Caballero & Tseng, 2016). Notably, social experience during juvenility has been found to be necessary for the development and function of fast-spiking, GABAergic parvalbumin (PV)-positive interneurons (Bicks et al., 2020). It is therefore likely that altered experience during early postnatal life may disrupt typical neural development, resulting in aberrant adult behavior. Evidence suggests that chronic adversity early in life affects the development of PFC PV interneurons (Schiavone et al., 2009). Both neonatal MS (Grassi-Oliveira et al., 2016; Holland et al., 2014) and juvenile SI (Bicks et al., 2020; Lukkes et al., 2012) have been found to decrease PFC PV levels. PV cells are preferentially enwrapped by perineuronal nets (PNNs; Enwright et al., 2016), specialized extracellular matrix structures that provide functional and physical protection, and regulation of cells that they enwrap (Cabungcal et al., 2013; Härtig et al., 1999). PFC PNN formation is protracted throughout development (Mauney et al., 2013), experiencing a spike in number during juvenility (Ueno et al., 2017) and maturing structurally until early adulthood (Gildawie et al., 2020). Compounding evidence indicates that delayed PNN maturation underlies critical period formation and susceptibility to adversity early in life (see review: Reichelt et al., 2019). Early life adversity impacts the development of PNNs in the PFC; animals exposed to post-weaning SI displayed reduced PNN intensity in the PFC (Ueno et al., 2017). It is also important to highlight the burgeoning exploration of sex-specific effects of adversity on PV+ PNNs. Neonatal MS was found to affect later-life PFC PV+ PNN structural integrity in a sex-dependent manner (Gildawie et al., 2020) and adolescent unpredictable chronic mild stress had sex- and age-specific effects on PNN/PV colocalization that correlated with increased anxiety-like behavior in females, but not males (Page & Coutellier, 2018). Together, these results implicate PNN/PV maturation in proper behavioral development, while the moderating effect of sex has yet to be fully understood.

Work in humans demonstrates that women exposed to multiple stressors during development have higher rates of depression and posttraumatic stress disorder in adolescence, compared to men (Suliman et al., 2009). Additionally, findings from human postmortem studies suggest that adults with affective disorders have fewer PFC PNNs, compared to controls (Alcaide et al., 2019; Mauney et al., 2013), suggesting that PNN aberrations may underly later-life susceptibility to neuropathology. Importantly, research suggests that multiple hits of adversity throughout the lifetime have additive effects on behavioral and GABAergic function later in life (Avital & Richter-Levin, 2005; Deslauriers et al., 2013; Monte et al., 2017). To our knowledge, no studies have investigated whether multiple hits of adversity have a synergistic effect on altered PFC PNN number and structural integrity. Additionally, it is unknown whether such changes occur specifically in female rats, mimicking increased vulnerability of young girls to multiple traumatic events and the subsequent onset of neuropsychiatric disorders. Here, we aimed to determine whether neonatal MS and juvenile SI results in female-specific compounding effects on adult PFC PNNs enwrapping PV-expressing interneurons that underlie susceptibility to anxiety-like behavior.

## 2. Materials and Methods

### 2.1. Subjects

All subjects were bred in an in-house colony with Sprague-Dawley rats originally obtained from Charles River Laboratories (Wilmington, MA). On P1, litters were culled to 10 pups (5 males and 5 females). Only one rat per litter was assigned to each experimental group to avoid potential litter effects. Animals were housed in standard polycarbonate wire-top cages with pine shaving bedding in a facility controlled for temperature (22-23°C) and humidity with a 12-hour light/dark cycle (light period 0700-1900, approximately 332 lux). Food (ProLab 5P00) and water (glass bottles) were available *ad libitum* to dams and weaned subjects. Experiments were carried out accordance with the 1996 Guide for the Care and Use of Laboratory Animals (NIH) with approval from the Institutional Animal Care and Use Committee at Northeastern University.

### 2.2. Developmental Adversity

#### 2.2.1. Maternal Separation

Beginning at postnatal day (P2), male and female pups were isolated from the dam and littermates for four hours per day (0900-1300) until P20, as previously described (Gildawie et al., 2020; Grassi-Oliveira et al., 2016; Holland et al., 2014; Honeycutt et al., 2020). Between P2 and P10, pups were separated in individual containers with bedding and droppings from the home cage – to assuage potential stress responses from altered olfactory cues – in a warm water bath (37°C). When pups were able to thermoregulate, separations were conducted in individual small cages containing home cage bedding (P11 to P20). All separations took place in a different room from where dams were located. Pups in control (Con) litters were handled for five minutes twice per week to alleviate handling disparities between groups due to separations. All litters underwent routine weekly husbandry and weighing on P9, P15, and P20 (approximately five min of handling).

#### 2.2.2. Juvenile Social Isolation

At weaning (P21), pups were separated from their dams and randomly assigned to be housed individually (SI) or pair-housed (PH) with a same-sex, condition-matched conspecific. Rats undergoing SI were housed in the same room as PH rats to have visual, auditory, and olfactory contact without any form of social interaction with littermates or conspecifics post-weaning (Möller et al., 2013b; Schiavone et al., 2009). At P35, all SI rats were housed with a condition- and sex-matched conspecific until experimentation.

### 2.3. Behavioral Testing

Rats were transported into a dimly lit testing room and left undisturbed to acclimate for 10-20 minutes. Anxiety-like behavior was tested in the elevated zero maze (EZM), a circular plexiglass alley elevated 70 cm off the ground. The alleyway was divided into four equal quadrants. Two of the quadrants (situated across the maze from one another) were walled with black plexiglass. The two remaining quadrants were unwalled and open to the surrounding testing environment. Each rat was placed in the closed area of the maze and left to explore the maze for five minutes. Maze exploration was recorded by an Ausdom HD 1080p Webcam and behavior was scored separately by an experimenter blind to experimental condition to determine time spent in open area (s), frequency to open, number of crossings, number of head pokes, and head poke duration (s). The maze was cleaned with 50% ethanol solution between each animal.

### 2.4. Immunohistochemistry

Fifteen days after behavioral testing (P85), animals were deeply anesthetized with CO2 and intracardially perfused with ice-cold 0.9% physiological saline followed by 4% paraformaldehyde (PFA) solution (see Fig. 1A for experimental timeline). Perfusions were performed between 0900 and 1200 to mitigate variability in PV and PNN expression throughout the day. After tissue collection, brains were post-fixed in PFA for three days and cryoprotected in 30% sucrose solution. Brains were sliced on a freezing microtome (Leica) to 40 μm sections and stored at −20°C in freezing solution until fluorescent staining.

**Figure 1.**
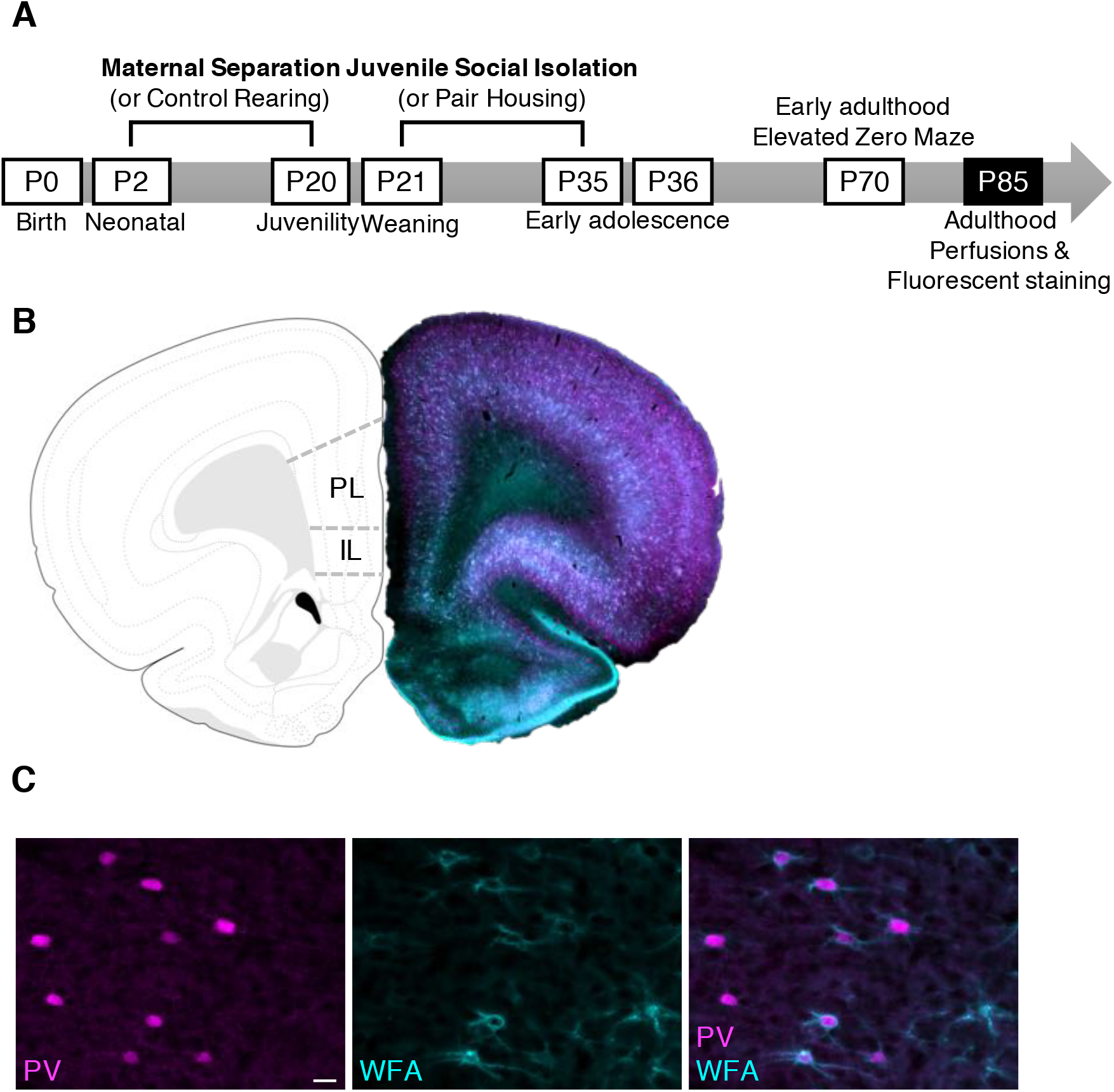
Experimental timeline and neuroanatomical representations. (A) Pups were exposed to maternal separation (or control rearing) postnatal day (P)2 to 20, followed by social isolation (or standard pair housing) P21-35. Rats were then pair housed until testing in the elevated zero maze on P70 and perfusions on P85. (B) Representative diagram (left) and stitched full-brain image (right) of the prelimbic (PL) and infralimbic (IL) prefrontal cortex. (C) Representative photomicrographs of parvalbumin (PV)-expressing neurons (magenta), *Wisteria floribunda* agglutinin (WFA)+ perineuronal nets (PNNs; cyan), and a merged image with WFA+ PNNs enwrapping non-PV cells (arrow), PV cells lacking PNNs (chevron), and PNNs surrounding PV cells (pentagonal arrow). Scale bar = 10 μm

Free-floating sections containing the PFC (between bregma 4.2 and 2.52 mm) were first washed in 1X phosphate buffered saline (PBS). The tissue was blocked in 5% normal donkey serum and 1% bovine serum albumin, then incubated in primary conjugate lectin from *Wisteria floribunda* agglutinin (WFA; 1:250, L1516, MilliporeSigma) and primary rabbit anti-PV antibody (1:1000, NB-120-11427, Novus Biologicals) at 4°C for 48 hours in PBS containing 0.3% Triton^™^ X-100 (PBST; Fisher Scientific). Sections were washed in PBST, followed by a three-hour incubation in secondary antibody solution composed of streptavidin conjugate 488 (1:3000, S32354, Thermofisher) and donkey anti-rabbit Alexa Fluor^®^ 350 (1:500, A10039, Invitrogen) in 0.3% PBST. Sections were washed in PBS and incubated in NeuroTrace^™^ 530/615 (1:200, N21482, Invitrogen) in PBS for one hour to visualize fluorescent Nissl staining. Sections underwent a final round of washes in PBS, then were mounted on positively-charged glass slides and coverslipped with ProLong Gold antifade mounting reagent (P36930, Invitrogen). Every round of staining included subjects from each experimental group to obviate batch-specific differences.

### 2.5. Microscopy and Image Quantification

WFA+ PNNs and PV+ interneurons were imaged using a Keyence BZ-X701 All-in-One fluorescent microscope. Z-stacks were captured at 20x magnification (image size: 724.69 μm x 543.52 μm) in three serial sections of the prelimbic (PL) and infralimbic (IL) cortices (see Fig. 1B for neuroanatomical diagram). The “Quick Full Focus” feature was used to yield one TIFF containing all images within the stack, encompassing all focal planes captured. Regions of interest were delineated using clearly visible landmarks and pre-defined boundaries according to a rat brain atlas (Paxinos & Watson, 2007) clearly visible via NeuroTrace^™^. Two sets of z-stacks were captured for the PL and one per IL, bilaterally (18 stacks per animal).

After image acquisition, each set of TIFFs was quantified as previously reported (Gildawie et al., 2020) using the “Perineuronal net Intensity Program for the Standardization and Quantification of ECM Analysis” (PIPSQUEAK; Slaker et al., 2016), a FIJI (Schindelin et al., 2012) macro plugin developed specifically to quantify PNNs and PV cells. The number and intensity of WFA+ PNNs, PV+ interneurons, and PNNs colocalized with PV were semi-automatically counted through the entirety of the stained tissue section (average 21 μm per section) by an experimenter blind to experimental condition. Images were first analyzed in PIPSQUEAK using predetermined parameters for the detection of WFA and PV staining. To better detect faint staining, the low background subtraction setting was used. An experimenter blind to experimental condition corrected for misidentification of PNNs and PV neurons to achieve the most accurate identification for each image. Double labeling of PNNs and PV neurons was then semi-automatically identified by PIPSQUEAK, with at least 80% overlap needed to be considered colocalized. A final approval step allowed the experimenter to correct any misidentified colocalization. Measures acquired were PV cell count and intensity, PNN count and intensity, colocalized PNN/PV count, intensity of PNNs surrounding PV cells, and intensity of PV cells surrounded by PNNs (see Fig. 1C for examples of PV and WFA staining).

### 2.6. Statistical Analyses

Statistical analyses were conducted using GraphPad Prism 7 software or IBM SPSS Statistics V.25. Prior to analysis, data were tested for outliers using Grubbs’ Test, homogeneity of variances was assessed with Levene’s Test of Equality of Error Variances, and Normality of residuals was evaluated via the Shapiro-Wilk Test of Normality (with assessment of skewness and kurtosis). Three-way ANOVAs were then performed to assess experimental effects of Sex, Rearing, and Housing in the EZM and immunohistochemical analysis. Main effects of Rearing or Housing, as well as interactions between Rearing and Housing or Sex and Rearing or Housing, were followed up with two-way ANOVAs (Rearing x Housing) separated by Sex to further assess the impact of multiple hits of developmental adversity on males and females. Effect size (η_p_^2^) was calculated in SPSS and categorized as small (η_p_^2^ = 0.01), medium (η_p_^2^ = 0.06), or large (ηp^2^ = 0.14; Richardson, 2011). Significant main effects and interactions were further assessed with Tukey’s HSD post-hoc tests to compare between groups, while correcting for multiple comparisons.

Pearson’s correlation analyses were run in Prism for each behavioral measure with significant group differences to determine the overall impact of behavior in the EZM on PFC structure in males and females separately. Further, we were interested in the effect of these Rearing and Housing conditions on the relationship between brain and behavior. We therefore used the “interactions” package (Long, 2019) in R (R Core Team, 2020) to perform multiple regression analyses with interactions. Regressions were run for each behavioral measure to assess the relationship between each PNN or PV measure (centered around the mean), Rearing condition, and Housing condition and anxiety-like behavior (independent variable). Models were run to assess two-way (Rearing x Brain and Housing x Brain) and three-way (Rearing x Housing x Brain) interactions. The unstandardized coefficient (B) is reported for each interaction regression. Each regression was assessed for deviations in Normality, homoskedasticity, and the presence of outliers. Given a significant two- or three-way interaction, follow-up regressions were run in Prism to assess the slope of each group’s regression line.

## 3. Results

### 3.1. Neonatal maternal separation and juvenile social isolation have opposing effects on adult risk assessment and hyperactivity in females

Anxiety-like behavior was measured in both males and females using the EZM following neonatal MS and juvenile SI. A three-way interaction between Rearing, Housing, and Sex was not apparent when the duration spent in the open area of the maze was assessed (*F*_1,82_ = 0.119, *p* = 0.731, η_p_^2^ = 0.001; Fig. 2A). There were also no significant main effects of Rearing (*F*_1,82_ = 0.380, *p* = 0.539, η_p_^2^ = 0.005), Housing (*F*_1,82_ = 3.101, *p* = 0.082, η_p_^2^ = 0.036), or Sex (*F*_1,82_ = 0.111, *p* = 0.740, η_p_^2^ = 0.001). Follow-up analyses were, therefore, not performed. As shown in Fig. 2B, however, the frequency of entrances to the open area was significantly impacted by juvenile SI (main effect of Housing: *F*_1,82_ = 6.505, *p* = 0.013, η_p_^2^ = 0.073) and was mildly affected by Rearing in a sex-dependent manner (Rearing x Sex interaction *F*_1,82_ = 4.126, *p* = 0.045, η_p_^2^ = 0.048). Follow-up two-way ANOVAs separated by Sex revealed that effects of Rearing (females: *F*_1,41_ = 7.085, *p* = 0.011, η_p_^2^ = 0.147; males: *F*_1,41_ = 0.164, *p* = 0.687, η_p_^2^ = 0.004) and Housing (females: *F*_1,41_ = 6.919, *p* = 0.012, η_p_^2^ = 0.144; males: *F*_1,41_ = 1.237, *p* = 0.273, η_p_^2^ = 0.029) were driven by females. Specifically, females exposed to MS entered the open area of the maze less than Con animals. Contrarily, SI females entered the open area *more* than PH counterparts. Tukey post-hoc comparisons show that females exposed to SI alone entered the open area significantly more than females exposed to MS alone (*p* = 0.0028). We also measured the number of times animals crossed from one enclosed area to the other, as a measure of locomotion (Fig. 2C). Similar to the frequency of entrances to the open area of the maze, there was a main effect of Housing (*F*_1,82_ = 8.576, *p* = 0.004, η_p_^2^ = 0.095) and an interaction between Rearing and Sex (*F*_1,82_ = 8.576, *p* = 0.007, η_p_^2^ = 0.086) on the number crossings from one closed area to the other. Again, when two-way ANOVAs were performed, females seemed to drive the decrease in crossings following MS (main effect of Rearing; females: *F*_1,41_ = 11.679, *p* = 0.001, η_p_^2^ = 0.222; males: *F*_1,41_ = 0.829, *p* = 0.368, η_p_^2^ = 0.020) and SI (main effect of Housing; females: *F*_1,41_ = 6.972, *p* = 0.012, η_p_^2^ = 0.145; males: *F*_1,41_ = 2.802, *p* = 0.102, η_p_^2^ = 0.064) exposure. Post-hoc analyses comparing each group revealed that the number of crossings was significantly higher in females exposed to only SI compared to those with only MS (*p* = 0.0005) or those exposed to both manipulations (Con, SI vs. MS, SI: *p* = 0.0110). These data demonstrate hyperactivity following juvenile SI. In contrast, MS showed an opposing effect, decreasing locomotion in adult females, but not males.

**Figure 2.**
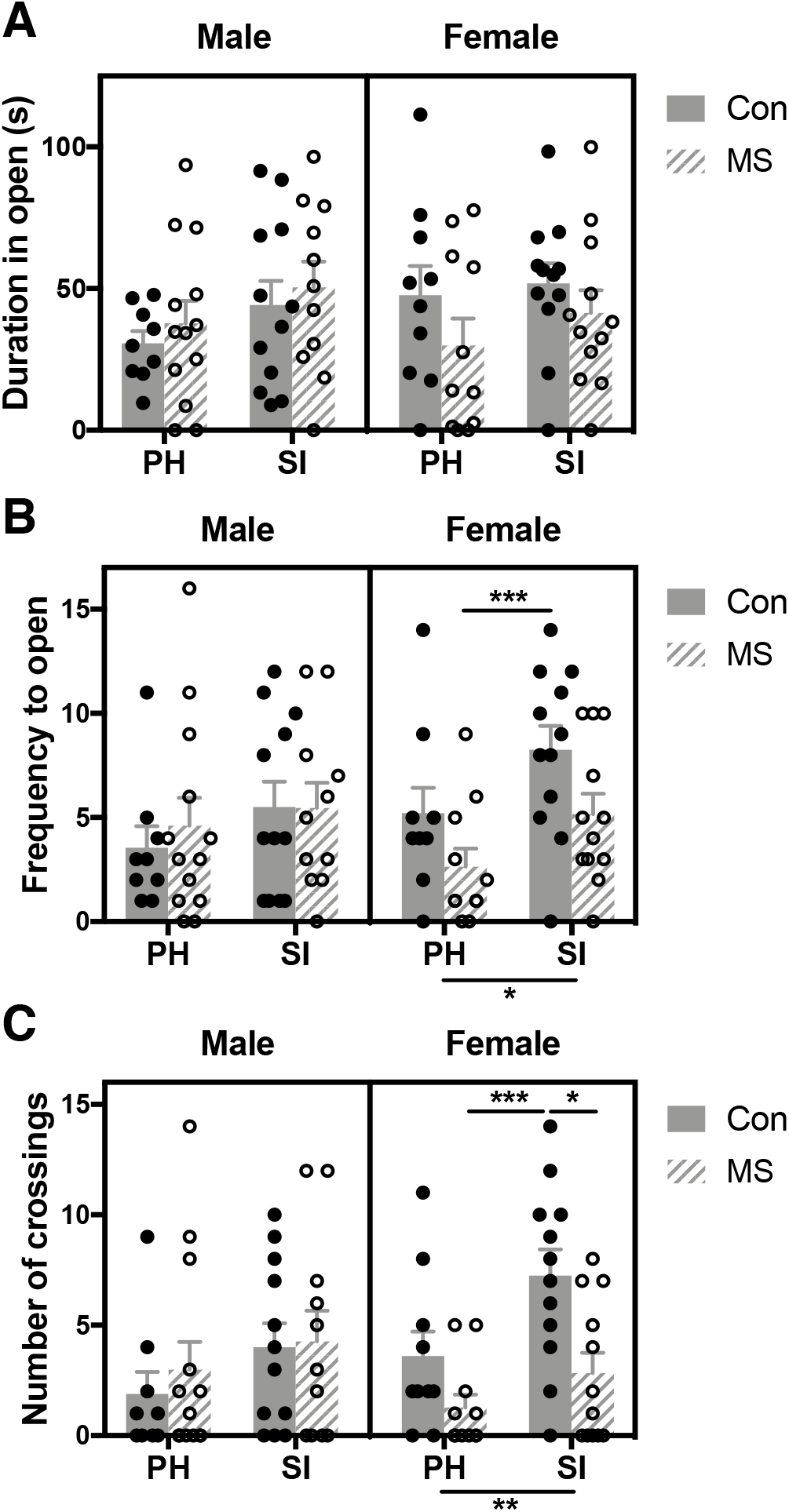
Effects of maternal separation (MS) and social isolation (SI) on adult anxiety-like behavior and locomotion in the elevated zero maze. (A) No effects of MS or SI on duration in the open area. (B) SI increased the frequency to the open area and (C) number of crossings in females that was opposed by MS. *: *p* < 0.05; **: *p* < 0.01; ***: *p* < 0.001; *n* = 9-13/group.

The number and duration of head pokes into the open area (without subsequent exit out of the closed area) was evaluated to gauge risk assessment, which has been previously shown to be a more sensitive measure of anxiety-like behavior (Roy & Chapillon, 2004; Sidor et al., 2010). As shown in Fig. 3A, we observed that the number of head pokes depended on Rearing, Housing, and Sex (*F*_1,82_ = 6.948, *p* = 0.010, η_p_^2^ = 0.078). Further analyses separated by Sex revealed that female (Rearing x Housing interaction *F*_1,41_ = 4.807, *p* = 0.034, η_p_^2^ = 0.105), but not male (*F*_1,41_ = 2.216, *p* = 0.144, η_p_^2^ = 0.051) head poke frequency depended on both Rearing and Housing. Follow-up post-hoc analysis revealed no group differences. When the duration of head pokes was assessed, a small – but significant – three-way interaction was observed (*F*_1,82_ = 4.390, *p* = 0.039, η_p_^2^ = 0.051). There was also an overall decrease following SI (main effect of Housing *F*_1,82_ = 4.590, *p* = 0.035, η_p_^2^ = 0.053) that was driven by a non-significant, but moderate decrease head poke duration in females (main effect of Housing; females: *F*_1,41_ = 4.026, *p* = 0.051, η_p_^2^ = 0.089; males: *F*_1,41_ = 2.416, *p* = 0.128, η_p_^2^ = 0.056; Fig. 3B). Follow-up two-way ANOVAs also showed a main effect of Rearing in females (*F*_1,41_ = 5.667, *p* = 0.022, η_p_^2^ = 0.121), but not males (*F*_1,41_ = 0.072, *p* = 0.789, η_p_^2^ = 0.002). Tukey post-hoc comparisons revealed that SI decreased the time spent head poking (MS, PH vs. Con, SI: *p* = 0.0158) and both hits of early life adversity prevented this decrease (Con, SI vs. MS, SI: *p* = 0.0400) in females. These findings suggest that – in females, but not males – juvenile SI results in decreased risk assessment behavior in females, while MS has the opposite effect.

**Figure 3.**
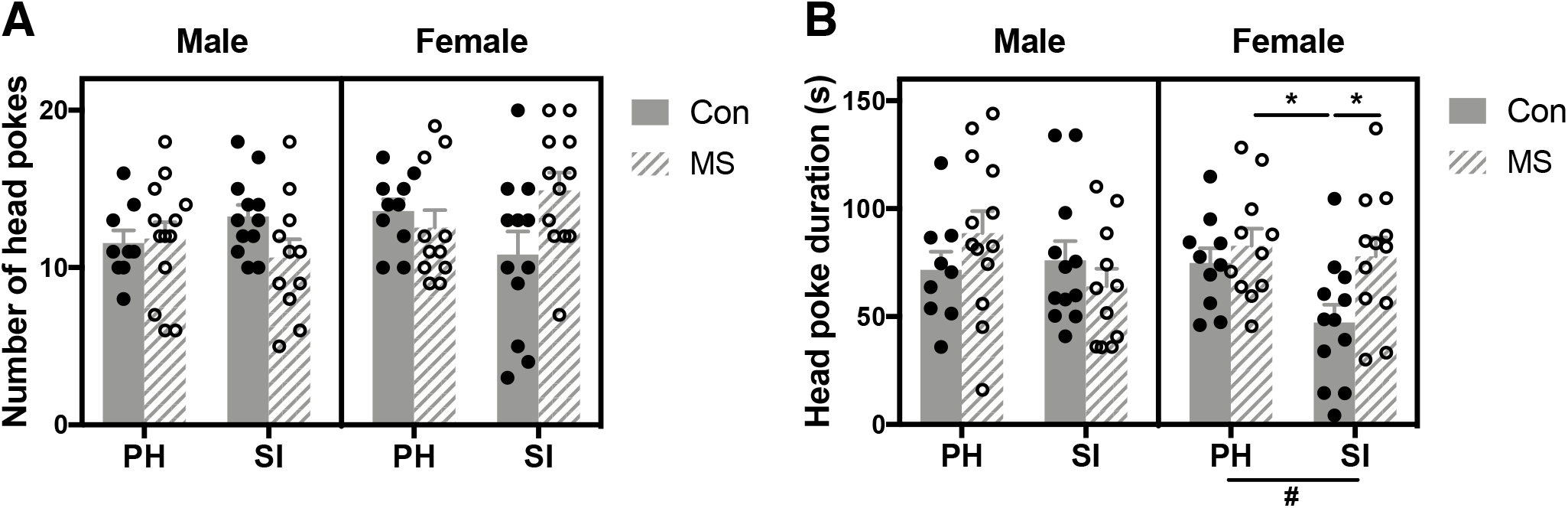
Effects of maternal separation (MS) and social isolation (SI) on adult risk assessment in the elevated zero maze. (A) The number of head pokes into – without entering – the open area depended on both MS and SI, which was driven by females. No group differences were observed. (B) Head poke duration was significantly decreased by SI in females with an opposing effect of MS. #: *p* < 0.1; *: *p* < 0.05; *n* = 9-13/group.

### 3.2. Early life adversity has compounding effects on female prefrontal cortex parvalbumin-expressing interneurons and perineuronal net structural integrity

#### 3.2.1. Prelimbic prefrontal cortex

##### Parvalbumin-expressing neuron density and intensity

The number and intensity of PV-expressing interneurons and the PNNs that enwrap them were assessed to determine whether there were sex-dependent effects of multiple adverse experiences throughout early development on the adult PFC. Three-way ANOVA revealed an interaction between Rearing, Housing, and Sex (*F*_1,75_ = 4.180, *p* = 0.044, η_p_^2^ = 0.053), as well as an overall decrease in PV count in the PL following MS (main effect of Rearing *F*_1,75_ = 7.409, *p* = 0.008, η_p_^2^ = 0.090; Fig. 4A). When males and females were analyzed separately, the effect of Rearing was found to be primarily driven by females (*F*_1,36_ = 4.191, *p* = 0.048, η_p_^2^ = 0.104), although males also demonstrated a nonsignificant, moderate decrease in PV neuron count (*F*_1,39_ = 3.165, *p* = 0.083, η_p_^2^ = 0.075). Tukey post-hoc comparisons reveal that females exposed to both MS and SI had fewer PV cells in the PL compared to females exposed to SI alone (*p* = 0.042), which was not apparent in males (*p* = 0.9542). When PV intensity was assessed, however, no effects were observed (Rearing x Housing x Sex interaction *F*_1,75_ = 1.513, *p* = 0.223, η_p_^2^ = 0.020; Fig. 4B).

**Figure 4.**
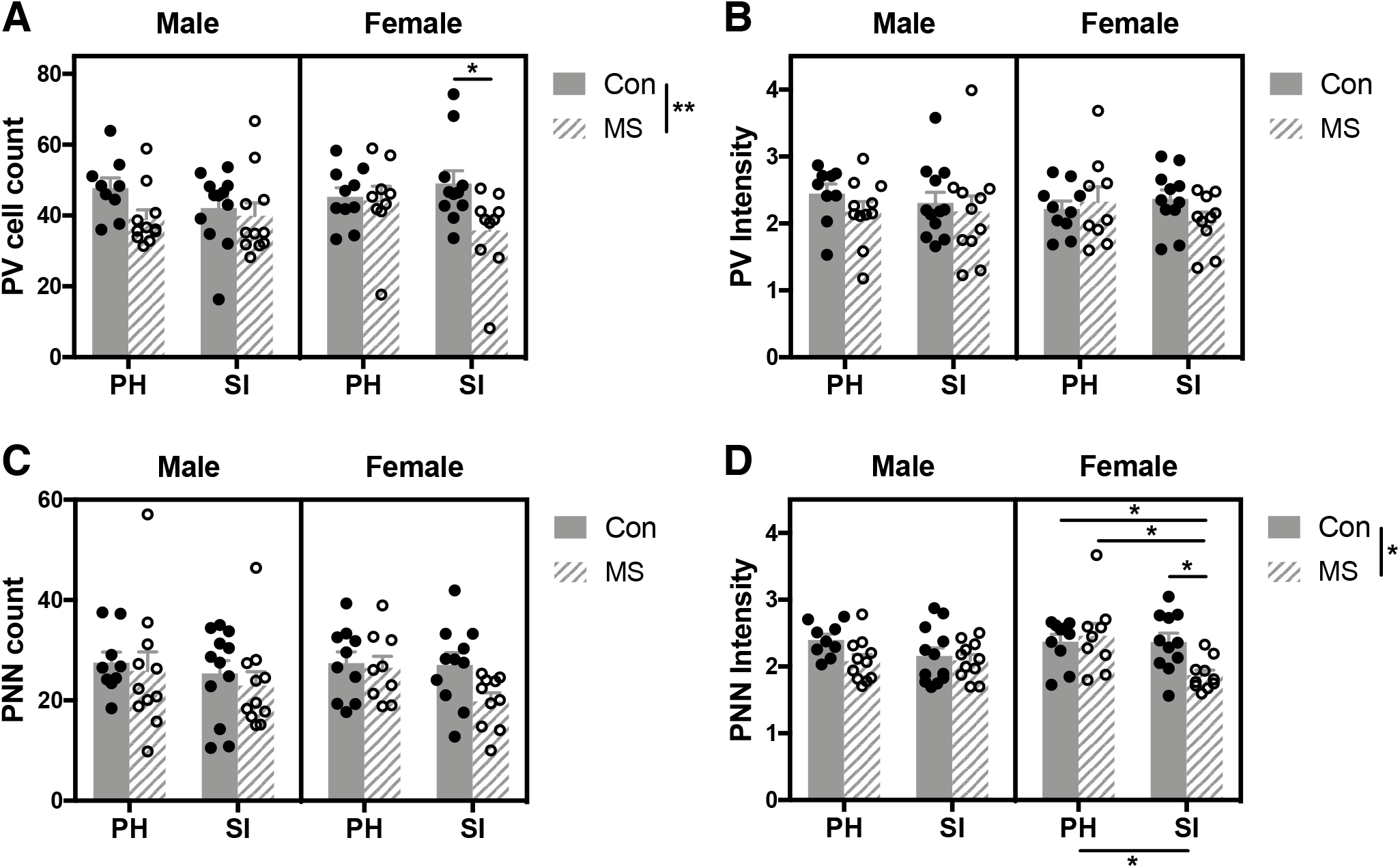
Effects of maternal separation (MS) paired with social isolation (SI) on adult parvalbumin (PV)-expressing interneurons and perineuronal nets (PNNs) in the prelimbic (PL) prefrontal cortex. (A) A three-way interaction between Rearing, Housing, and Sex was revealed, where females exposed to both MS and SI had fewer PV cells in the PL than females exposed to SI alone. (B) Intensity of PV immunofluorescence and (C) PNN number were not altered. (D) An additive effect of MS and SI was apparent in female PNN intensity, where females exposed to MS and SI had PNNs that fluoresced less than all other female groups. *: *p* < 0.05; **: *p* < 0.01; *n* = 9-13/group.

##### Perineuronal net density and intensity

We also quantified the number and intensity of PNNs following MS and SI. While Rearing, Housing, and Sex did not change the number of PL PNNs (*F*_1,75_ = 0.560, *p* = 0.457, η_p_^2^ = 0.007; Fig. 4C), PNN intensity was significantly altered (Rearing x Housing x Sex interaction *F*_1,75_ = 6.722, *p* = 0.011, η_p_^2^ = 0.082; Fig. 4D). Both MS (main effect of Rearing *F*_1,75_ = 5.940, *p* = 0.017, η_p_^2^ = 0.073) and SI (main effect of Housing *F*_1,75_ = 6.434, *p* = 0.013, η_p_^2^ = 0.079) decreased overall PNN intensity. Separate two-way ANOVAs in males and females revealed an interaction between Rearing and Housing (*F*_1,36_ = 5.214, *p* = 0.028, η_p_^2^ = 0.127) and a main effect of Housing (*F*_1,36_ = 5.312, *p* = 0.027, η_p_^2^ = 0.129) in females. Meanwhile, males showed no Rearing x Housing interaction (*F*_1,39_ = 1.593, *p* = 0.214, η_p_^2^ = 0.039) and a modest, nonsignificant decrease in PNN intensity following MS (main effect of Rearing *F*_1,39_ = 0.3456, *p* = 0.071, η_p_^2^ = 0.081), suggesting that the overall Rearing x Housing interaction was driven by females. Moreover, Tukey post-hoc comparisons show that PNN intensity in females exposed to both MS and SI was significantly lower than Con, SI (*p* = 0.0365); MS, PH (*p* = 0.0159); and Con, PH (*p* = 0.0410) females. These findings demonstrate an additive effect of MS and SI on PL PNN intensity, exclusively in females.

###### Colocalized parvalbumin and perineuronal net density and intensity

Analysis of the number of PL PNNs specifically enwrapping PV neurons revealed an overall decrease following MS (main effect of Rearing *F*_1,75_ = 7.632 *p* = 0.007, η_p_^2^ = 0.092), as well as a three-way interaction (*F*_1,75_ = 5.085, *p* = 0.027, η_p_^2^ = 0.063; Fig. 5A). Two-way ANOVAs split by Sex revealed a significant interaction between Rearing and Housing in females (*F*_1,36_ = 4.503, *p* = 0.041, η_p_^2^ = 0.111), but not males (*F*_1,39_ = 1.116, *p* = 0.297, η_p_^2^ = 0.028). Both males and females showed a moderate, but nonsignificant MS-induced decrease in PNN+ PV cell count (main effect of Rearing; females: *F*_1,36_ = 4.012, *p* = 0.053, η_p_^2^ = 0.10; males: *F*_1,39_ = 3.627, *p* = 0.064, η_p_^2^ = 0.085). Planned assessment of group differences showed that females exposed to both MS and SI had significantly fewer PV neurons surrounded by PNNs, compared to females exposed to SI alone (*p* = 0.0245). This difference was not apparent in males (*p* = 0.9240). The proportion of PV cells surrounded by PNNs, however, did not change depending on differences in Rearing, Housing, or Sex (*F*_1,75_ = 0.621, *p* = 0.443, η_p_^2^ = 0.008; Fig. 5B). Analysis of intensity measures for colocalized PV cells and PNNs revealed that the intensity of PV neurons surrounded by PNNs was also unaltered (*F*_1,75_ = 2.118, *p* = 0.150, η_p_^2^ = 0.027; Fig. 5C). No main effects (Rearing *F*_1,75_ = 1.597, *p* = 0.220, η_p_^2^ = 0.021; Housing *F*_1,75_ = 0.268, *p* = 0.606, η_p_^2^ = 0.004) were apparent. When the intensity of PNNs enwrapping PV neurons was assessed, a small three-way interaction was observed (*F*_1,75_ = 4.567, *p* = 0.036, η_p_^2^ = 0.057). Additionally, MS (main effect of Housing *F*_1,75_ = 5.001, *p* = 0.028, η_p_^2^ = 0.063) and SI (main effect of Housing *F*_1,75_ = 7.331, *p* = 0.008, η_p_^2^ = 0.089) significantly decreased PV+ PNN intensity (Fig. 5D). Follow-up analysis in males and females separately showed that the effect of Housing (females: *F*_1,36_ = 5.639, *p* = 0.023, η_p_^2^ = 0.135; males: *F*_1,39_ = 1.70, *p* = 0.20, η_p_^2^ = 0.042) was driven by females. Additionally, an interaction between Rearing and Housing was non-significant, but moderate in females (*F*_1,36_ = 3.784, *p* = 0.060, η_p_^2^ = 0.095), with no effect in males (*F*_1,39_ = 0.873, *p* = 0.356, η_p_^2^ = 0.022). Multiple comparisons showed that PNNs surrounding PV cells in animals exposed to both MS and SI fluoresced significantly less than those that underwent MS alone (females: *p* = 0.0252; males: *p* = 0.9933) and Con Rearing and PH (females: *p* = 0.0410; males: *p* = 0.2040) in females, but not males.

**Figure 5.**
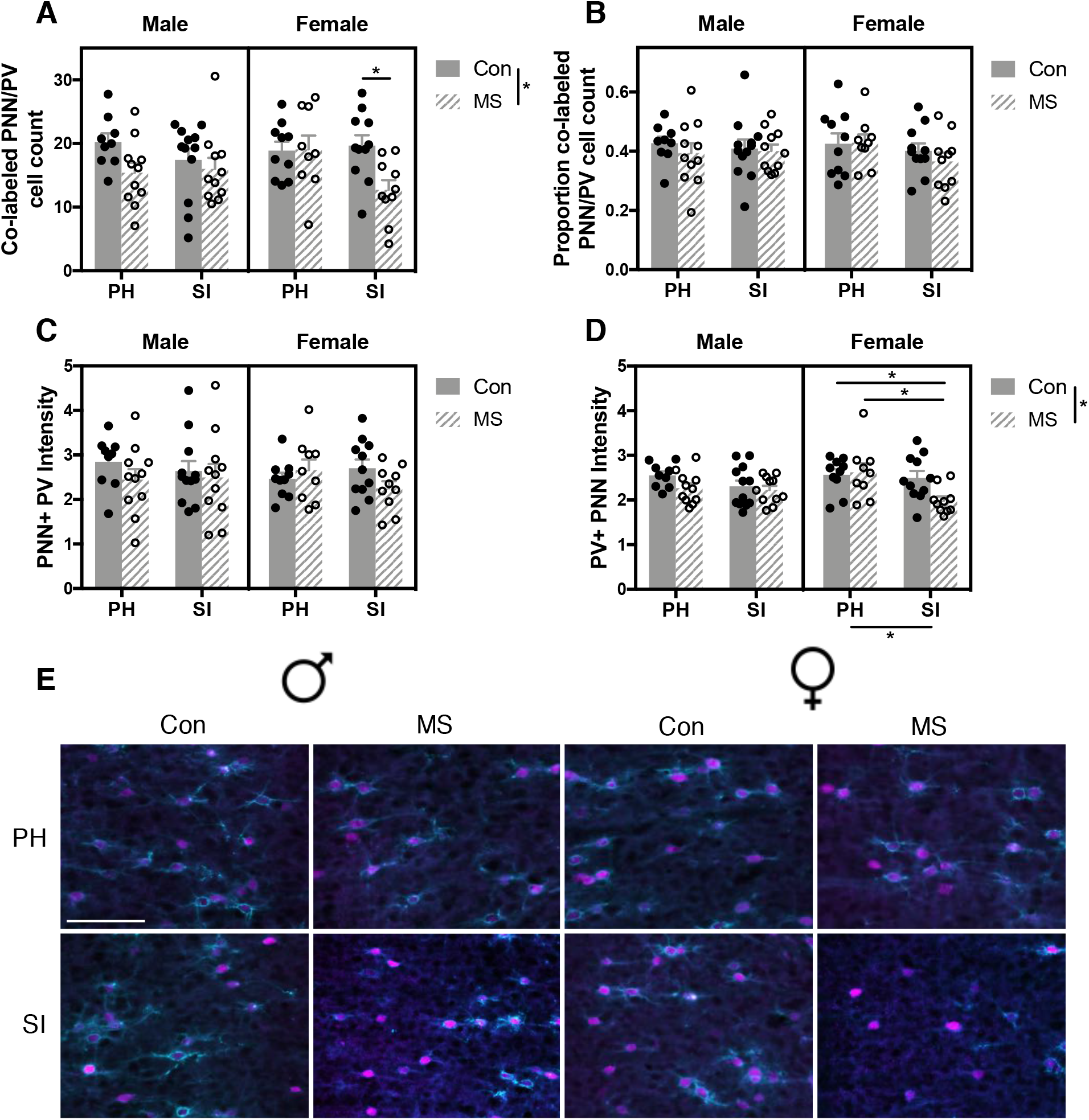
Effects of maternal separation (MS) paired with social isolation (SI) on co-labeled parvalbumin (PV)-expressing neurons and perineuronal nets (PNNs) in the prelimbic (PL) prefrontal cortex. (A) The number of PNN+ PV cells was decreased in both males and females in the PL, but only females showed a significant reduction in PNN+ PV number following both MS and SI exposure. (B) Neither the proportion nor (C) intensity of PV cells enwrapped by PNNs was affected by Rearing or Housing condition. (D) An additive effect of MS and SI on the intensity of PNNs surrounding PV cells in the PL was observed only females. (E) Representative photomicrographs of adult *Wisteria floribunda* agglutinin (WFA; cyan)+ PNNs and PV cells (magenta) in the PL. Scale bar = 50 μm; *: *p* < 0.05; *n* = 9-13/group

#### 3.2.2. Infralimbic prefrontal cortex

##### Parvalbumin-expressing neuron density and intensity

To investigate potential region-specific effects of multiple adverse experiences, we also assessed density and intensity measures in the IL. Three-way ANOVA revealed that PV density was decreased by MS (main effect of Rearing *F*_1,75_ = 9.475, *p* = 0.003, η_p_^2^ = 0.112), but no interaction between Rearing, Housing, and Sex was observed (*F*_1,75_ = 0.386, *p* = 0.536, η_p_^2^ = 0.005; Fig. 6A). Follow-up twoway ANOVAs revealed a decrease in PV number following MS in both females (main effect of Rearing *F*_1,36_ = 5.799, *p* = 0.021, η_p_^2^ = 0.139) and males (*F*_1,39_ = 3.816, *p* = 0.058, η_p_^2^ = 0.089), although the effect in males was non-significant, but moderate. Multiple comparisons revealed no group differences. When we assessed PV intensity in the IL, no three-way interaction was observed (*F*_1,75_ = 0.584, *p* = 0.447, η_p_^2^ = 0.008; Fig. 6B).

**Figure 6.**
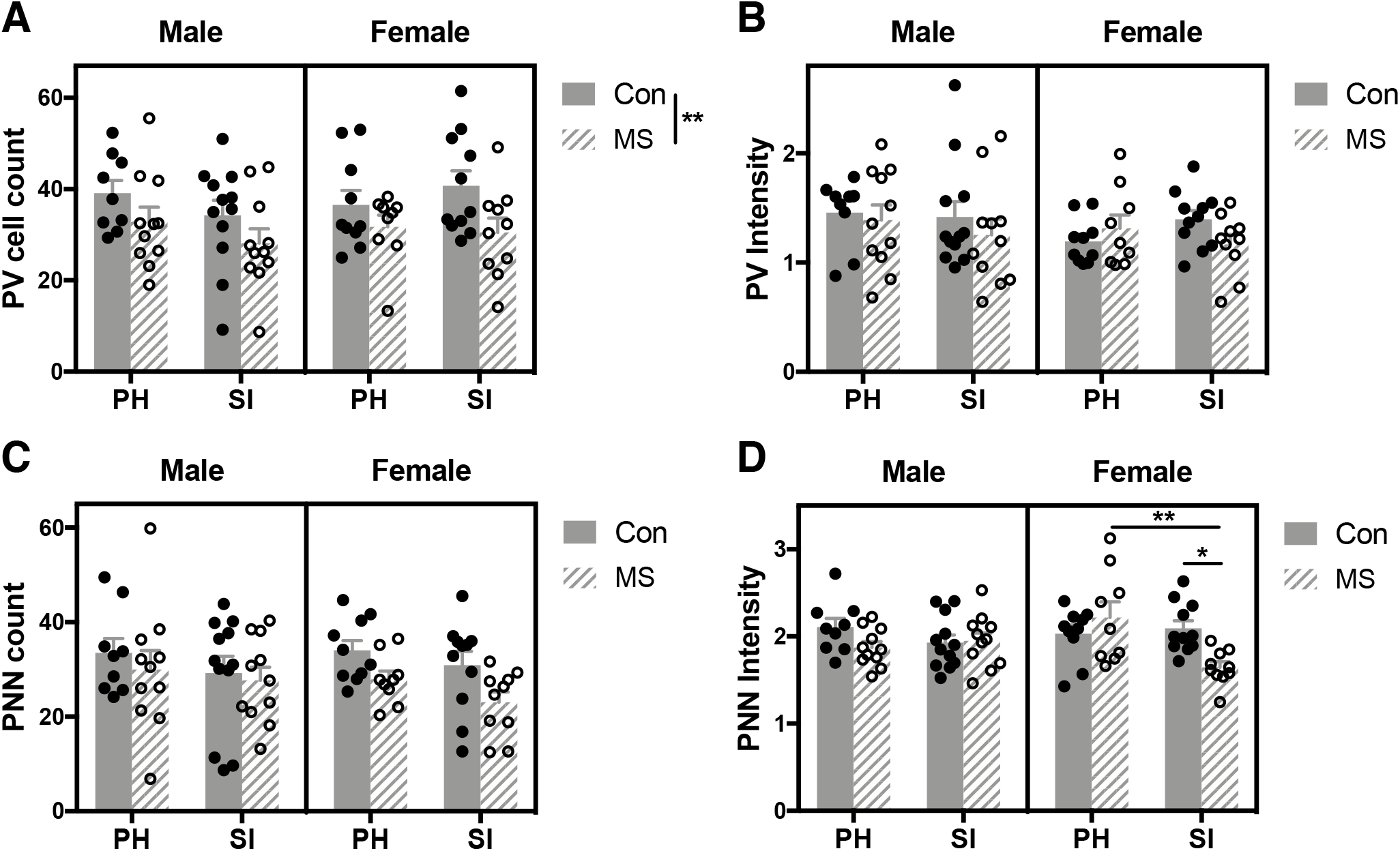
Effects of maternal separation (MS) paired with social isolation (SI) on adult parvalbumin (PV)-expressing interneurons and perineuronal nets (PNNs) in the infralimbic (IL) prefrontal cortex. (A) PV cell count was decreased in both males and females. (B) Intensity of PV immunofluorescence and (C) PNN number were not changed by Rearing or Housing. (D) A three-way interaction between Rearing, Housing, and Sex was revealed, where an additive effect of MS and SI occurred in females only. *: *p* < 0.05; **: *p* < 0.01; *n* = 9-13/group.

##### Perineuronal net density and intensity

Further analysis of PNNs in the IL showed no alterations in PNN number due to Rearing, Housing, and Sex (*F*_1,75_ = 0.203, *p* = 0.653, η_p_^2^ = 0.003; Fig. 6C); however, a three-way interaction was observed when PNN intensity was assessed (*F*_1,75_ = 10.237, *p* = 0.002, η_p_^2^ = 0.120; Fig. 6D). Follow-up two-way ANOVAs separated by Sex showed a strong interaction between Rearing and Housing (females: *F*_1,36_ = 8.364, *p* = 0.006, η_p_^2^ = 0.189; males: *F*_1,39_ = 2.142, *p* = 0.151, η_p_^2^ = 0.052), as well as decrease in PNN intensity following SI (main effect of Housing; females: *F*_1,36_ = 5.467, *p* = 0.025, η_p_^2^ = 0.132; males: *F*_1,39_ = 0.345, *p* = 0.560, η_p_^2^ = 0.009) only in females. Tukey post-hoc comparisons revealed that females exposed to both MS and SI showed lower fluorescent intensity than females exposed to either MS (*p* = 0.0049) or SI (*p* = 0.0267) alone. These group differences were not apparent in males (*p* = 0.9217; *p* = 0.9979).

##### Colocalized parvalbumin and perineuronal net density and intensity

MS significantly decreased the overall number of PV neurons enwrapped by PNNs in the IL (main effect of Rearing *F*_1,75_ = 9.775, *p* = 0.003, η_p_^2^ = 0.115); however, there was no three-way interaction between Rearing, Housing, and Sex (*F*_1,75_ = 0.942, *p* = 0.335, η_p_^2^ = 0.012; Fig. 7A). Follow-up two-way ANOVAs showed a main effect of Rearing in females (*F*_1,36_ = 7.861, *p* = 0.008, η_p_^2^ = 0.179), with a non-significant, but moderate, effect in males (*F*_1,39_ = 2.956, *p* = 0.093, η_p_^2^ = 0.070). Tukey post-hoc comparisons revealed no group differences, however. The proportion (*F*_1,75_ = 0.912, *p* = 0.343, η_p_^2^ = 0.012; Fig. 7B) and intensity (*F*_1,75_ = 0.645, *p* = 0.424, η_p_^2^ = 0.009; Fig. 7C) of PV cells enwrapped by PNNs were not altered by Rearing, Housing, or Sex. When the intensity of PNNs surrounding PV neurons was assessed, a three-way interaction was observed (*F*_1,75_ = 7.671, *p* = 0.007, η_p_^2^ = 0.093) and SI decreased PV+ PNN intensity (*F*_1,75_ = 5.416, *p* = 0.023, η_p_^2^ = 0.067; Fig. 7D). Separate two-way ANOVAs demonstrate an SI-induced decrease in PV+ PNN intensity (main effect of Housing; females: *F*_1,36_ = 4.178, *p* = 0.048, η_p_^2^ = 0.104; males: *F*_1,40_ = 1.201, *p* = 0.280, η_p_^2^ = 0.030), exclusively in females. Additionally, there was a significant interaction between Rearing and Housing in females (*F*_1,36_ = 6.292, *p* = 0.017, η_p_^2^ = 0.149), but not males (*F*_1,40_ = 1.432, *p* = 0.239, η_p_^2^ = 0.035). Analysis of group differences showed that females – but not males – exposed to MS and SI had PNNs enwrapping PV cells that fluoresced significantly less than those exposed to SI alone (females: *p* = 0.0419; males: *p* = 0.9562) and MS alone females: *p* = 0.0169; males: *p* = 0.9999), suggesting a sex-specific additive effect of multiple hits of adversity throughout development.

**Figure 7.**
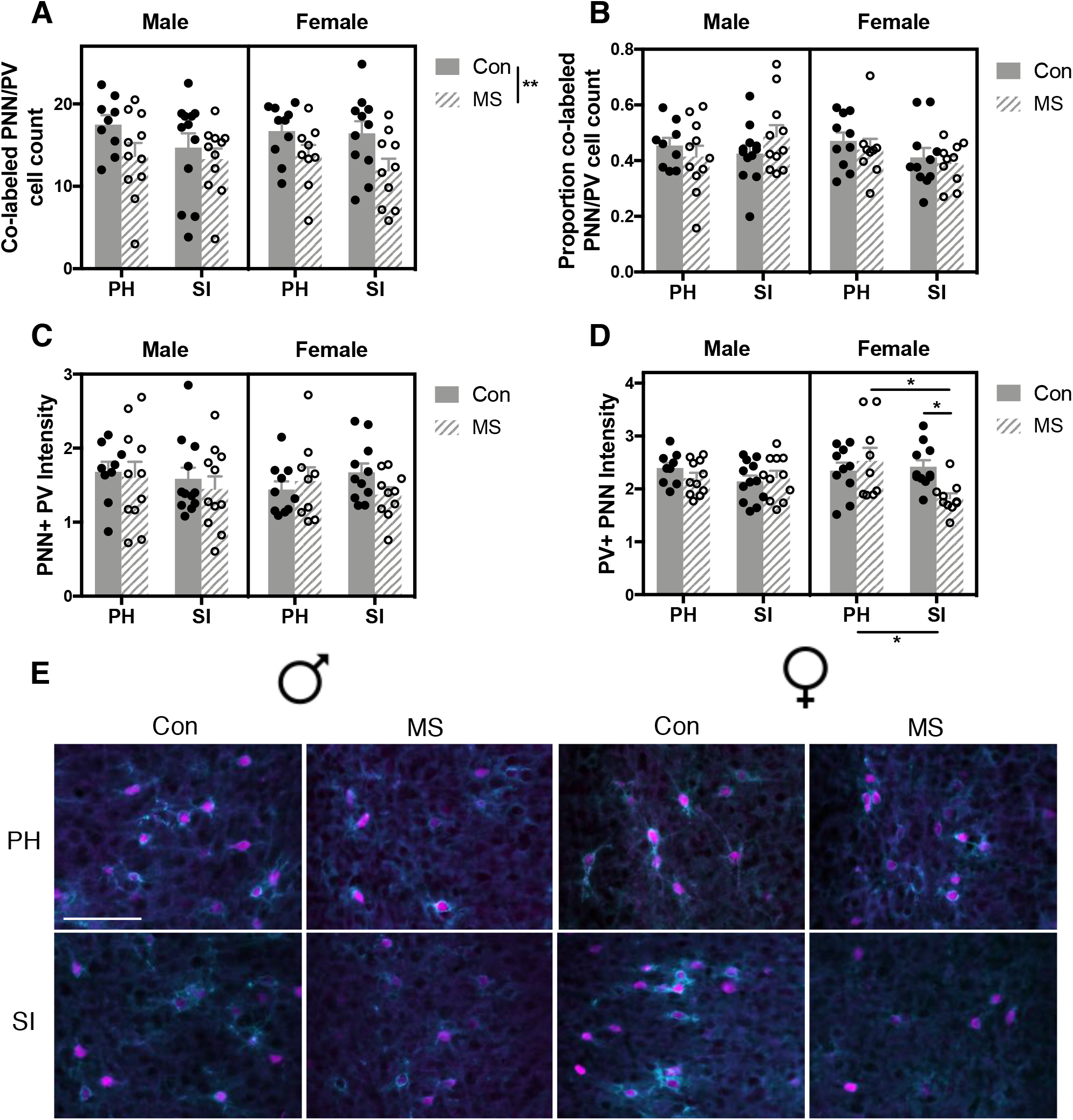
Effects of maternal separation (MS) paired with social isolation (SI) on co-labeled parvalbumin (PV)-expressing neurons and perineuronal nets (PNNs) in the infralimbic (IL) prefrontal cortex. (A) The number of PNN+ PV cells was decreased in the IL of both males and females. (B) The proportion and (C) intensity of PV cells surrounded by PNNs were not altered. (D) An additive effect of MS and SI on the intensity of PNNs surrounding PV cells in the PL was observed only in females. (E) Representative photomicrographs of adult *Wisteria floribunda* agglutinin (WFA; cyan)+ PNNs and PV cells (magenta) in the PL. Scale bar = 50 μm; *: *p* < 0.05; *n* = 9-13/group

### 3.3. Relationship between anxiety-like behavior and prefrontal cortex neural structure differs in males and females

To determine the relationship between PFC cellular measures and performance in the EZM, Pearson’s correlation analyses were conducted on behavioral measures that were significantly impacted by Rearing or Housing, with PV and PNN measures (see Tables 1 and 2 for correlations in the PL and IL, respectively). There was an inverse correlation between the intensity of PV neurons enwrapped by PNNs in the PL and frequency to the open area of the maze in males (*r* = −0.3124, *p* = 0.0440), but not females (*r* = −0.0712, *p* = 0.6626). This correlation was not apparent in IL cells (males: *r* = −0.2281, *p* = 0.1462; females: *r* = 0.0127, *p* = 0.9380). The number of PV cells in the PL was correlated inversely with the number of crossings in males (*r* = −0.3306, *p* = 0.0305) and positively in females (*r* = 0.3183, *p* = 0.0453). Again, this relationship did not extend to the IL (males: *r* = −0.2781, *p* = 0.0745; females: *r* = 0.2607, *p* = 0.1042). The intensity of PL PV neurons surrounded by PNNs was also inversely correlated with the number of crossings in males (*r* = −0.3689, *p* = 0.0162), but not females (*r* = −0.0571, *p* = 0.7264). This correlation was not significant in the IL (males: *r* = −0.2625, *p* = 0.0931); females: *r* = −0.0014, *p* = 0.9932). There was also positive correlation between PNN+ PV intensity and head poke duration in males in both the PL (*r* = 0.3252, *p* = 0.0356) and IL (*r* = 0.3049, *p* = 0.0496). No significant correlation was found in females (PL: *r* = 0.0947, *p* = 0.5612; IL: *r* = - 0.0492, *p* = 0.7628).

**Table 1.** Pearson’s correlation analyses between measures of prelimbic parvalbumin (PV) cell count, co-labeled perineuronal net (PNN)/PV cell count, PNN+ PV intensity, and PV+ PNN intensity with frequency to open, number of crossings, and head poke duration in the elevated zero maze. *: *p* < 0.05 (also indicated in bold).

|  | Frequency to open |  |
| --- | --- | --- |
|  | Male | Female |
| PV count | $r = -0.2445, p = 0.1186$ | $r = 0.3027, p = 0.0576$ |
| PNN+ PV count | $r = -0.1148, p = 0.4690$ | $r = 0.2677, p = 0.0949$ |
| PNN+ PV intensity | **$r = -0.3124, p = 0.0440^*$** | $r = -0.0712, p = 0.6626$ |
| PV+ PNN intensity | $r = -0.0067, p = 0.9663$ | $r = 0.1291, p = 0.4274$ |
|  | Number of crossings |  |
|  | Male | Female |
| PV count | **$r = -0.3306, p = 0.0305^*$** | **$r = 0.3183, p = 0.0453^*$** |
| PNN+ PV count | $r = -0.1829, p = 0.2463$ | $r = 0.2524, p = 0.1161$ |
| PNN+ PV intensity | **$r = -0.3689, p = 0.0162^*$** | $r = -0.0571, p = 0.7264$ |
| PV+ PNN intensity | $r = -0.0669, p = 0.6737$ | $r = 0.1760, p = 0.2773$ |
|  | Head poke duration |  |
|  | Male | Female |
| PV count | $r = 0.1914, p = 0.2246$ | $r = -0.1399, p = 0.3891$ |
| PNN+ PV count | $r = 0.0344, p = 0.8287$ | $r = -0.1280, p = 0.4311$ |
| PNN+ PV intensity | **$r = 0.3252, p = 0.0356^*$** | $r = 0.0947, p = 0.5612$ |
| PV+ PNN intensity | $r = 0.0326, p = 0.8379$ | $r = -0.1538, p = 0.3247$ |

**Table 2.** Pearson’s correlation analyses between measures of infralimbic parvalbumin (PV) cell count, co-labeled perineuronal net (PNN)/PV cell count, PNN+ PV intensity, and PV+ PNN intensity with frequency to open, number of crossings, and head poke duration in the elevated zero maze. *: *p* < 0.05 (also indicated in bold).

|  | Frequency to open |  |
| --- | --- | --- |
|  | Male | Female |
| PV count | $r = -0.1855, p = 0.2396$ | $r = 0.2059, p = 0.2024$ |
| PNN+ PV count | $r = -0.0064, p = 0.9680$ | $r = 0.2433, p = 0.1304$ |
| PNN+ PV intensity | $r = -0.2281, p = 0.1462$ | $r = 0.0127, p = 0.9380$ |
| PV+ PNN intensity | $r = 0.0517, p = 0.7452$ | $r = 0.0910, p = 0.5764$ |
|  | Number of crossings |  |
|  | Male | Female |
| PV count | $r = -0.2781, p = 0.0745$ | $r = 0.2607, p = 0.1042$ |
| PNN+ PV count | $r = -0.0712, p = 0.6540$ | $r = 0.2945, p = 0.0651$ |
| PNN+ PV intensity | $r = -0.2625, p = 0.0931$ | $r = -0.0014, p = 0.9932$ |
| PV+ PNN intensity | $r = -0.1208, p = 0.4765$ | $r = 0.1036, p = 0.5247$ |
|  | Head poke duration |  |
|  | Male | Female |
| PV count | $r = 0.0983, p = 0.5357$ | $r = -0.1283, p = 0.4300$ |
| PNN+ PV count | $r = -0.1187, p = 0.4540$ | $r = -0.0873, p = 0.5922$ |
| PNN+ PV intensity | **$r = 0.3049, p = 0.0496^*$** | $r = -0.0492, p = 0.7628$ |
| PV+ PNN intensity | $r = -0.0060, p = 0.9699$ | $r = -0.0173, p = 0.9156$ |

### 3.4. Rearing and Housing affect the relationship between brain and behavior differently in males and females

#### 3.4.1. Prelimbic prefrontal cortex

Linear regressions with interactions were performed separately in males and females to determine whether correlations (or lack thereof) between brain and behavior were due to group-specific moderation of the overall relationship, (see Supp Table 1 for results from all PL regressions). When the number of crossings was assessed, there was significant interaction between Housing and the intensity of PL PNNs surrounding PV cells in females (B = 5.1866, *p* = 0.0261; Fig. 8B), but not males (B = 1.5853, *p* = 0.6725; Fig. 8A), suggesting that Housing condition altered the relationship between the number of crossings and PV+ PNN intensity. A Housing x Brain interaction was found in males (B = 1.9563, *p* = 0.0474; Fig. 8C) when the relationship between overall PV number and duration of head pokes was assessed. This interaction was not apparent in females (B = −0.8261, *p* = 0.3596; Fig. 8D). Similarly, Housing altered the relationship between the number of co-labeled PNNs and PV cells and head poke duration in males (B = 4.172, *p* = 0.0213; Fig. 8E), but not females (B = −1.0562, *p* = 0.5442; Fig. 8F). Follow-up analyses showed no significantly non-zero slopes when groups were separated (data not shown).

**Figure 8.**
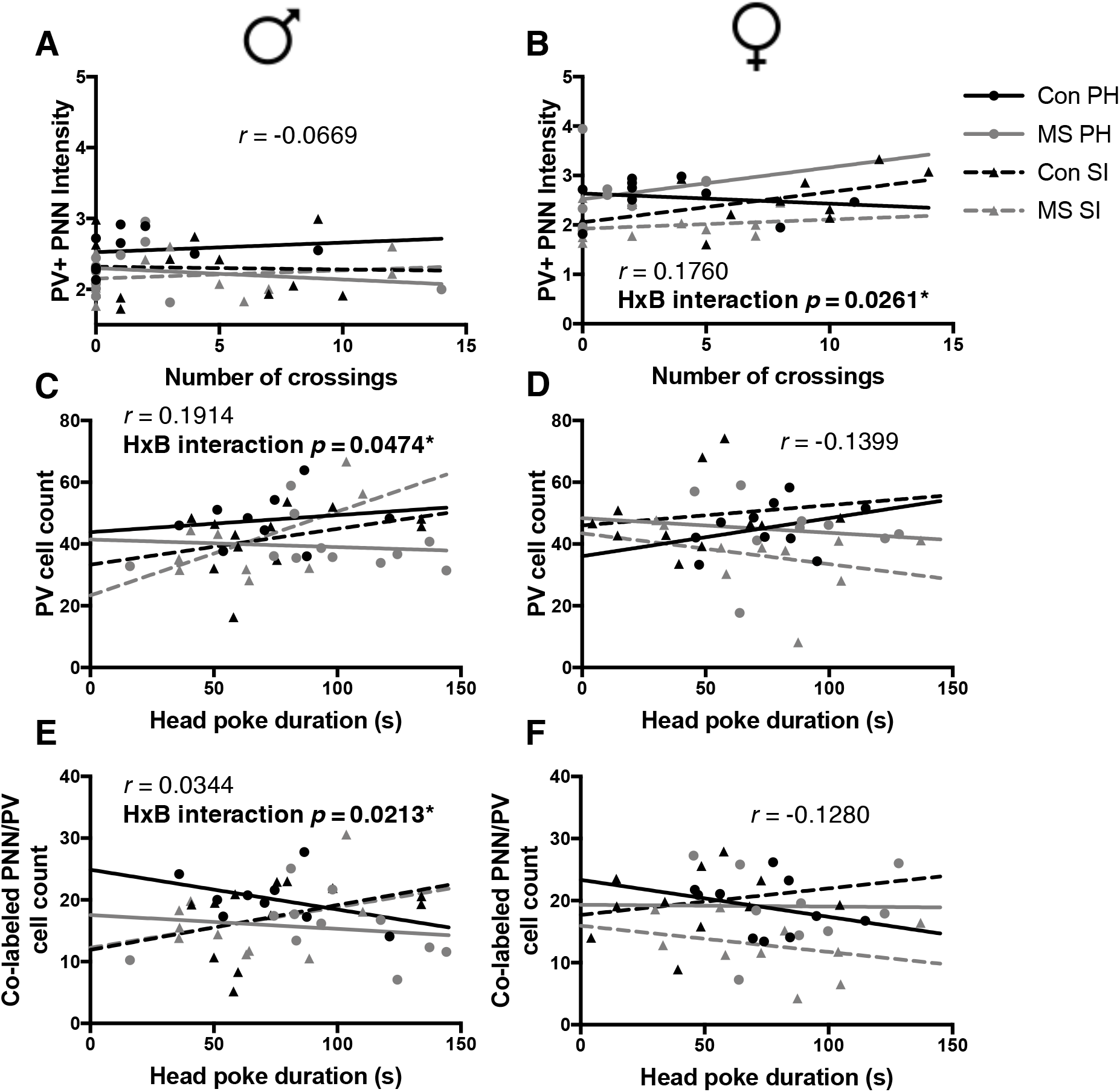
Two-and three-way interaction regression analyses between measures of prelimbic parvalbumin (PV) cell count, co-labeled perineuronal net (PNN)/PV cell count, PNN+ PV intensity, and PV+ PNN intensity with frequency to open, number of crossings, and head poke duration in the elevated zero maze. Dependent variables for these regressions were Rearing (R), Housing (H), and PNN/PV measure – labeled Brain (B) for simplicity. The Pearson’s correlation coefficient (*r*) from the overall correlation analysis and the *p*-value corresponding to the unstandardized regression coefficient from the interaction analysis are reported. *: *p* < 0.05 (also indicated in bold).

#### 3.4.2. Infralimbic prefrontal cortex

In the IL (see Supp Table 2 for results from all IL regressions), there was a significant three-way interaction (Rearing x Housing x Brain) when assessing the number of crossings and intensity of PNNs surrounding PV neurons in females (B = −13.180, *p* = 0.0215; Fig. 9B), suggesting that both Rearing and Housing mediate the relationship between crossings and PV+ PNN intensity. This interaction was moderate, but not significant in males (B = 15.6009, *p* = 0.0544; Fig. 9A). When the duration of head pokes was assessed, an interaction was revealed between Housing and PV cell count in males (B = 2.1413, *p* = 0.0245; Fig. 9C), but not females (B = −1.6558, *p* = 0.0925; Fig. 9D). A slight interaction between Housing and the number of colabeled PNNs and PV cells on head poke duration, only in males (males: B = 3.938, *p* = 0.0475; Fig. 9E; females: B = −3.1310, *p* = 0.1746; Fig. 9F). Lastly, a three-way interaction was apparent in males (B = −130.877, *p* = 0.0282; Fig. 9G) – but not females (B = 44.732, *p* = 0.3469; Fig. 9H) – on the relationship between head poke duration and the intensity of PNNs surrounding PV cells. Similar to the PL, no slopes were significantly different from zero (data not shown).

**Figure 9.**
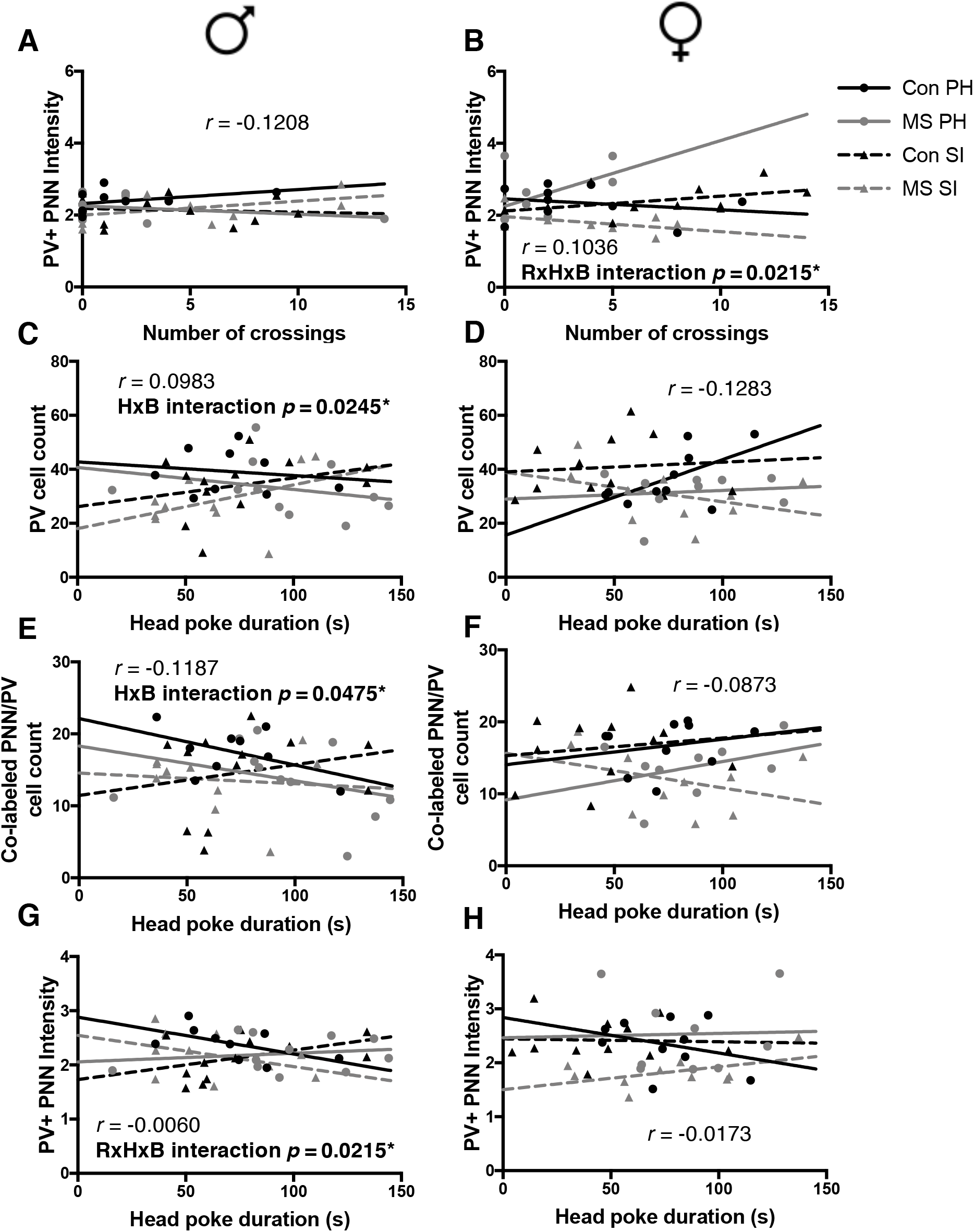
Two-and three-way interaction regression analyses between measures of infralimbic parvalbumin (PV) cell count, co-labeled perineuronal net (PNN)/PV cell count, PNN+ PV intensity, and PV+ PNN intensity with frequency to open, number of crossings, and head poke duration in the elevated zero maze. Dependent variables for these regressions were Rearing (R), Housing (H), and PNN/PV measure – labeled Brain (B) for simplicity. The Pearson’s correlation coefficient (*r*) from the overall correlation analysis and the *p*-value corresponding to the unstandardized regression coefficient from the interaction analysis are reported. *: *p* < 0.05 (also indicated in bold).

## 4. Discussion

A vast literature of clinical and preclinical research provides evidence that adversity throughout early development has long-term consequences in the brain that impact behavior (see reviews: Andersen & Teicher, 2008; Ganguly & Brenhouse, 2015; Heim & Binder, 2012; Pechtel & Pizzagalli, 2011). Multiple episodes of such adversity throughout childhood have been further associated with increased neuropsychiatric symptoms, particularly in women (Manyema et al., 2018; Suliman et al., 2009). However, there is a paucity of plausible biological mechanisms to explain sex-specific disruptions to affect-regulating brain circuitry following one or more early life stressors. In the current study, we modeled adversity during two windows of susceptibility (neonatal and juvenile stages) in the form of MS and SI to determine whether there is a compounding effect of multiple hits of adversity on adult anxiety-like behavior, as well as PV-expressing interneurons and WFA-labeled PNNs in the PFC. Our findings suggest a sex-specific impact of early life adversity on adult locomotion and risk-assessment behavior, where females are more sensitive to adversity-induced deviations in performance. Our data also indicate a sexspecific additive effect of adversity on PNN structural integrity and PV cell count. Additionally, we find that manipulating the developmental environment impacts the relationship between PFC neural structure and anxiety-like behavior differently in males and females.

Behaviorally, our data indicate no change in the amount of time spent in the open area of the EZM, signifying a lack of effect of repeated developmental adversity on this measure of anxiety-like behavior. We observed an increase in frequency to the open area in SI females, which may be driven by increased locomotion, displayed by crossings from one closed area of the EZM to the other. Contrary to our hypothesis, this finding may suggest a SI-induced decrease in anxiety-like behavior. In past work, adolescent SI (P28-70) has been found to have similar anxiolytic effects in female C57BL/6J mice, as well as hypersociality (Rivera-Irizarry et al., 2020). While social behavior was not measured in the current study, it is possible that the observed hyperactivity may be explained by increased motivation for social contact. Notably, SI females also showed decreased risk-assessment behavior, suggested in the NIH Research Domain Criteria (National Institute of Mental Health, 2016) to reflect vigilance to potential threats. MS, however, had a contrasting effect on EZM performance, resulting in an overall decrease in frequency to open and crossings that was driven by females, suggesting that females may experience long term behavioral effects of neonatal adversity. Females exposed to MS paired with SI showed a decrease in locomotion and an increase in risk-assessment behavior compared to females exposed to SI alone. This may suggest that the SI-induced increase in exploratory behavior and reduced risk assessment is prevented by MS experience. We have previously shown that MS leads to increased anxiety-liked behavior in adolescent males (Ganguly et al., 2015) and others have found that MS paired with early weaning resulted in increased anxiety-like behavior in adult males (Murthy et al., 2019). It is essential, however, to recognize the limitations of behavioral tasks – such as the EZM and Elevated Plus Maze – as indicators of anxiety-like behavior. Rodent-based anxiety-related tasks rely on observable behavior. In humans, however, Generalized Anxiety Disorder does not commonly manifest as distinct behavioral phenotypes, instead existing as a more internal, psychophysiological experience (Troisi, 1999). Importantly, meta-analyses have found a large amount of heterogeneity and discrepancy from studies assessing behavioral effects of early life adversity, especially between different measures of anxiety-like behavior (Bonapersona et al., 2019; Daniel Wang et al., 2020). Such conceptual and procedural concerns complicate generalization of rodent behavioral paradigms to human experience (Wall & Messier, 2001). Clinical literature points to aberrations in childhood and adolescent environment on gender disparities in later-life depression (Heim et al., 2009; Leach et al., 2008) and PTSD (McLaughlin et al., 2010). Findings regarding gender differences in anxiety following childhood adversity, however, are mixed (Gallo et al., 2018). While our findings may point to a role of early life adversity in female-specific behavioral development, future work in humans and animal models must disentangle the complex relationship between childhood maltreatment and mental health in males and females.

Few studies have investigated the effect of multiple hits of adversity on later-life PV interneurons and PNNs, and the current work is the first to assess these in the context of sex differences. Here, we demonstrate that neonatal MS paired with juvenile SI decreases the number of PV-expressing interneurons in the PFC and reduces the structural integrity of the PNNs that enwrap them, in females only. Previously, Castillo-Gomez and colleagues (2017) did not observe a potentiated decrease in PFC PV cell count, PNN number, or PV+ PNN number following two hits of stress in male mice, corroborating our finding that compounding neurostructural deficits may occur specifically in females. The decrease in PV cell number seen in females exposed to both MS and SI did not occur in tandem with a decrease in PNN number, suggesting that the potentiated reduction in PNN intensity did not occur simply because there were fewer PV cells in the PFC. The lack of decrease in the proportion of PNN coverage, however, demonstrates that the overall PV decrease was not driven solely by the population of PV cells enwrapped by PNNs. Our data confirm prior evidence that both populations of PV cells (PNN-lacking and PNN-enwrapped) are vulnerable to the effects of stress (Cabungcal et al., 2013). Further, PV cells enwrapped by structurally aberrant PNNs may be at similar risk to those lacking any protection from PNNs. PNNs act as a physical and molecular barrier, with many extracellular components working in concert to maintain structural integrity and protect the cells that they enwrap (Wang & Fawcett, 2012). We have previously found that MS alone delays PNN formation in the PFC with normal levels apparent by adulthood (Gildawie et al., 2020), suggesting a transient effect of early life adversity on altered PNN expression. While altered PNN structural integrity in juvenility may have long term consequences on brain maturation, the current findings suggest that a second hit of juvenile adversity is required for prolonged PNN reduction in adulthood. Importantly, we have also found that PV cells in the PFC of juveniles exposed to neonatal MS undergo increased oxidative damage (Soares et al., 2020). As degradation of PNNs via chondroitinase ABC has been found to increase PV neuron vulnerability to oxidative stress (Cabungcal et al., 2013), this converging evidence suggests that a lack of PNNs following MS contributes to oxidative damage and consequential PV loss upon subsequent challenges.

PV neuron maturation during early development is essential for long term function. Reduction in adolescent PFC PV expression has been shown to decrease GABAergic transmission in adulthood, altering excitatory-inhibitory synaptic activity (Caballero et al., 2020). Indeed, PV reduction has long been implicated in mood disorders, such as schizophrenia (Beasley & Reynolds, 1997; Enwright et al., 2016). Importantly, the presence of PNNs has an important impact on synaptic input and physiology of PV positive cells (Riga et al., 2017). Typical PNN number and integrity is therefore critical for maintaining proper neural transmission throughout life. In postmortem human studies, individuals with schizophrenia (Mauney et al., 2013) and bipolar disorder (Alcaide et al., 2019) expressed fewer PNNs in the PFC, suggesting that PV and PNN disruption has important implications for the onset of neuropsychiatric disorders. These studies, however, were not sufficiently powered to assess differences between men and women. Our findings suggest that female PNNs and PV cells in the PFC are more susceptible to the effects of repeated adversity, which may underlie the increased risk of developing mood disorders for young girls following multiple instances of adversity during childhood. While changes in PV cell function were not assessed in the current work, a decrease in PFC PNN structural integrity following adversity during key developmental stages could have important consequences for cortical plasticity with implications for sex-specific response to stress.

Proper GABAergic function is critical for cognitive and affective development (see review: Cristo, 2007). We therefore assessed the relationship between PV/PNN number and structural integrity on performance in the EZM. Interestingly, no regressions that showed significant two- or three-way interactions had coinciding significant correlations. This may suggest an impact of adversity on the relationship between brain and behavior that plays a role in neutralizing the overall correlations. When we assessed the relationship between PV and PNN measures and locomotion-related behaviors (frequency to open and number of crossings), we found that males overall had more correlations between PV count and intensity measures and behavior than females, despite showing no group differences in solely behavioral analysis. Interestingly, males with fewer PV neurons in the PL – but not IL – showed increased locomotion, while the opposite was true for females, suggesting a potential sex difference in the relationship between locomotion and PV expression. Further, we found that the relationship between brain and locomotion-related behavior was not modulated by Housing in males, but was in females. When assessing the relationship between the brain and risk-assessment behavior, males also showed a positive correlation between PNN+ PV intensity and head poke duration. Unlike locomotion, however, the relationship between PV/PNN measures and risk-assessment behavior was not modulated by Housing condition in females, but was in males. Taken together, these findings suggest sex-specific relationships between brain, behavior, and adversity in males and females that differ according to behavioral subtype. While scarce, evidence is beginning to point to sexspecific neural mechanisms underlying similar behavioral phenotypes (Becker & Chartoff, 2019). It is possible that males and females regulate anxiety-like behavior through separate mechanisms; however, more work is needed to identify these mechanisms and what this means for gender-specific therapeutic mediation following childhood adversity. Additionally, while there were no substantial differences in PV or PNN expression between the PL and IL, behavior was less correlated with PNN/PV expression in the IL versus the PL. Evidence suggests that the PL and IL play different roles in the expression of anxiety-like behavior, where PL activation results in an anxiogenic response and IL activation does not (Suzuki et al., 2016). Such region-specific differences in behavioral mediation may underlie the disparate relationships between brain and behavior in the current work.

When interpreting the current work, some considerations should be made. First, in designing the current study, we sought to quantify the relationship between adversity-induced PNN/PV disruption and anxiety-like behavior; however, behavioral and neuroanatomical assessment was separated by 15 days, making interpretation of the brain-behavior relationship more complex. Since PNNs and PV cells are both highly sensitive to altered experience, even in adulthood (Riga et al., 2017), we chose to include a gap between behavioral testing and tissue collection to avoid potential confounds from the task. Secondly, we chose to use Pipsqueak for cell count and intensity analyses. Pipsqueak is a powerful tool that can provide faster and more reliable quantification than more manual methods (Slaker et al., 2016). We are, however, limited in our analysis, as Pipsqueak – in its current form – does not allow for count or intensity quantification for PNNs surrounding non-PV cells. Prior work from our lab has shown distinct differences in how MS affects the expression of PV cells with and without PNNs (Gildawie et al., 2020). Given the important role of PNNs in PV expression and function, however, measuring the structural integrity of PNNs enwrapping PV cells versus other cell types could provide important information regarding cell-specific effects of early life adversity. Lastly, in adult mice previously exposed to MS, both males and females demonstrated enhanced anxiety-like behavior; however, the effects in females were moderated by estrous phase (Romeo et al., 2003). While the current study was not powered to assess estrous phase, it is possible that female behavioral response to developmental adversity may be modulated by cycle. That said, female behavior was no more variable than that of males.

## 5. Conclusions

Early life is rife with sensitive periods during which aberrant experience can have lasting effects on plasticity and behavioral development (Bicks et al., 2020; Chocyk et al., 2010). Heightened neural malleability during early stages of development may – in part – stem from protracted formation of PNNs (Gildawie et al., 2020; Mauney et al., 2013), and may also depend on sex and pubertal status (Drzewiecki et al., 2020). Here, we show a clear sex-specific effect of multiple hits of early life adversity during distinct windows of susceptibility, where females demonstrate a potentiated decrease in PV number and PNN structural integrity following neonatal MS and juvenile SI. Evidence suggests that reduced PNN number and structural integrity may result in a detrimental increase in plasticity (see review: Reichelt et al., 2019) and expose enwrapped PV-expressing interneurons to the effects of oxidative stress (Cabungcal et al., 2013). PNN degradation has been seen to disrupt GABAergic circuitry (Riga et al., 2017) and hamper synaptic transmission (Blosa et al., 2015), which has implications for behavioral dysfunction (Bicks et al., 2020). In the current study, we observe a sex-specific effect of adversity on hyperactivity and risk-assessment behavior, as well as moderating effects of adversity on the brain-behavior relationship.

Recent work has elucidated potential mechanisms by which PNNs are degraded in response to altered experience and neuropsychiatric conditions. For example, microglia have been implicated in PNN structural development and degradation (Crapser et al., 2020; Crapser et al., 2020; Nguyen et al., 2020). Further, sex differences have been observed in the neuroimmune response to stress (Fonken et al., 2018; Gildawie & Orso et al., 2020), suggesting a microglia-mediated mechanism may underlie differences between males and females in their behavioral and neural response to repeated adversity during sensitive periods of development. The evidence presented here has important implications for sex-specific neurostructural and behavioral effects of multiple hits of adversity. Moreover, the current work helps lay the foundation for mechanistic investigation of sex-specific changes in plasticity, which will further our understanding of neuropsychiatric sequalae and potential preventative therapies.

## Abbreviations

PFC: prefrontal cortex
PL: prelimbic
IL: infralimbic
PNN: perineuronal net
WFA: *Wisteria floribunda* agglutinin
MS: maternal separation
Con: control
SI: social isolation
PH: pair-housed
P: postnatal day
PBS: phosphate buffered saline
PFA: paraformaldehyde
EZM: elevated zero maze

## Acknowledgements

We would like to thank Julia Terry, Nicholas Craffey, Michael Meding, Tobias Kremsmayer, Jason Hirsch, and Habiba Shaheed for their technical assistance in the preparation of this manuscript.

