## Supplementary Tables 1 & 2 for "A two-hit adversity model in developing rats reveals sex-specific impacts on prefrontal cortex structure and behavior"

|  | Frequency to open | | | | | |
| --- | --- | --- | --- | --- | --- | --- |
|  | Male | | | Female | | |
|  | Rearing  x Brain | Housing  x Brain | R x H x Brain | Rearing  x Brain | Housing  x Brain | R x H x Brain |
| PV count | B = 0.0131 *p* = 0.920 | B = -0.0509 *p* = 0.6859 | B = -0.2487 *p* = 0.3890 | B = -0.0494 *p* = 0.6682 | B = 0.0465 *p* = 0.6734 | B = -0.350 *p* = 0.1513 |
| PNN+ PV count | B = 0.0183 *p* = 0.9408 | B = -0.0732 *p* = 0.7553 | B = -0.1364 *p* = 0.8080 | B = -0.0996 *p* = 0.6705 | B = 0.0712 *p* = 0.7373 | B = -0.30 *p* = 0.5430 |
| PNN+ PV intensity | B = 0.1813 *p* = 0.9117 | B = -0.7075 *p* = 0.6531 | B = 1.1338 *p* = 0.7536 | B = 1.2780 *p* = 0.5630 | B = -0.0895 *p* = 0.9675 | B = 0.0501 *p* = 0.9917 |
| PV+ PNN intensity | B = 0.6511 *p* = 0.8619 | B = 0.4952 *p* = 0.8896 | B = 6.8661 *p* = 0.4276 | B = -2.5122 *p* = 0.3377 | B = 3.9792 *p* = 0.1122 | B = -1.7644 *p* = 0.7791 |
|  | Crossings | | | | | |
|  | Male | | | Female | | |
|  | Rearing  x Brain | Housing  x Brain | R x H x Brain | Rearing  x Brain | Housing  x Brain | R x H x Brain |
| PV count | B = -0.0942 *p* = 0.4746 | B = -0.0968 *p* = 0.4364 | B = -0.3302 *p* = 0.2307 | B = -0.1258 *p* = 0.2424 | B = 0.1171 *p* = 0.2640 | B = -0.2382 *p* = 0.2922 |
| PNN+ PV count | B = -0.2178 *p* = 0.4012 | B = -0.0794 *p* = 0.7423 | B = -0.3538 *p* = 0.5250 | B = -0.1690 *p* = 0.4443 | B = 0.2578 *p* = 0.2060 | B = -0.34734 *p* = 0.4518 |
| PNN+ PV intensity | B = -0.6175 *p* = 0.7195 | B = -0.5039 *p* = 0.7540 | B = 0.2433 *p* = 0.9467 | B = 0.1521 *p* = 0.9418 | B = 1.3296 *p* = 0.5302 | B = 1.6033 *p* = 0.7217 |
| PV+ PNN intensity | B = 0.6851 *p* = 0.8646 | B = 1.5853 *p* = 0.6725 | B = 8.4455 *p* = 0.3446 | B = -2.1679 *p* = 0.3783 | **B = 5.1866 *p* = 0.0261*** | B = -4.5249 *p* = 0.4228 |
|  | Head poke duration | | | | | |
|  | Male | | | Female | | |
|  | Rearing  x Brain | Housing  x Brain | R x H x Brain | Rearing  x Brain | Housing  x Brain | R x H x Brain |
| PV count | B = -0.0234 *p* = 0.9818 | **B = 1.9563 *p* = 0.0474*** | B = 1.3348 *p* = 0.5299 | B = -0.621 *p* = 0.5046 | B = -0.8261 *p* = 0.3596 | B = -0.1159 *p* = 0.9516 |
| PNN+ PV count | B = -0.6457 *p* = 0.7410 | **B = 4.172 *p* = 0.0213*** | B = -1.850 *p* = 0.6502 | B = -0.2429 *p* = 0.8940 | B = -1.0562 *p* = 0.5442 | B = -4.709 *p* = 0.2260 |
| PNN+ PV intensity | B = 8.4890 *p* = 0.5032 | B = 12.2630 *p* = 0.3223 | B = 10.9622 *p* = 0.4210 | B = -16.608 *p* = 0.3213 | B = -7.053 *p* = 0.6798 | B = -25.016 *p* = 0.4810 |
| PV+ PNN intensity | B = 9.140 *p* = 0.7558 | B = 30.604 *p* = 0.2770 | B = -95.415 *p* = 0.1386 | B = 16.388 *p* = 0.4126 | B = -19.5680 *p* = 0.3376 | B = -31.69 *p* = 0.5186 |

**Supp Table 1.** Two-and three-way interaction regression analyses between measures of prelimbic parvalbumin (PV) cell count, co-labeled perineuronal net (PNN)/PV cell count, PNN+ PV intensity, and PV+ PNN intensity with frequency to open, number of crossings, and head poke duration in the elevated zero maze. Dependent variables for these regressions were Rearing (R), Housing (H), and PNN/PV measure – labeled Brain for simplicity. The unstandardized regression coefficient (B) with the corresponding *p*-value are reported for each regression. *: *p* < 0.05 (also indicated in bold).

|  | Frequency to open | | | | | |
| --- | --- | --- | --- | --- | --- | --- |
|  | Male | | | Female | | |
|  | Rearing  x Brain | Housing  x Brain | R x H x Brain | Rearing  x Brain | Housing  x Brain | R x H x Brain |
| PV count | B = -0.0093 *p* = 0.9412 | B = -0.1031 *p* = 0.3989 | B = -0.3542 *p* = 0.2006 | B = 0.1406 *p* = 0.3054 | B = 0.1217 *p* = 0.3356 | B = -0.2625 *p* = 0.3370 |
| PNN+ PV count | B = 0.0629 *p* = 0.8172 | B = -0.0070 *p* = 0.9786 | B = -0.3614 *p* = 0.5795 | B = 0.0390 *p* = 0.9013 | B = 0.2011 *p* = 0.4839 | B = -0.3211 *p* = 0.6205 |
| PNN+ PV intensity | B = -0.0233 *p* = 0.9925 | B = 0.1351 *p* = 0.9538 | B = 2.5253 *p* = 0.6443 | B = 2.0452 *p* = 0.5229 | B = -3.7471 *p* = 0.2311 | B = -8.793 *p* = 0.1757 |
| PV+ PNN intensity | B = 2.2653 *p* = 0.5396 | B = 0.8242 *p* = 0.8247 | B = 12.9069 *p* = 0.1074 | B = -2.3101 *p* = 0.3685 | B = 2.0512 *p* = 0.4005 | B = -10.6462 *p* = 0.0921 |
|  | Crossings | | | | | |
|  | Male | | | Female | | |
|  | Rearing  x Brain | Housing  x Brain | R x H x Brain | Rearing  x Brain | Housing  x Brain | R x H x Brain |
| PV count | B = -0.0756 *p* = 0.5633 | B = -0.1098 *p* = 0.3793 | B = -0.3750 *p* = 0.180 | B = 0.0494 *p* = 0.7027 | B = 0.1692 *p* = 0.1564 | B = -0.1668 *p* = 0.5135 |
| PNN+ PV count | B = -0.1230 *p* = 0.6724 | B = 0.0709 *p* = 0.7947 | B = -0.3627 *p* = 0.5897 | B = -0.0231 *p* = 0.9375 | B = 0.3086 *p* = 0.2529 | B = -0.3606 *p* = 0.5417 |
| PNN+ PV intensity | B = -0.6174 *p* = 0.8166 | B = 1.0035 *p* = 0.6808 | B = 1.7925 *p* = 0.7511 | B = -0.08174 *p* = 0.7863 | B = -0.8463 *p* = 0.7813 | B = -7.480 *p* = 0.2311 |
| PV+ PNN intensity | B = 3.0314 *p* = 0.4436 | B = 3.6735 *p* = 0.3431 | B = 15.6009 *p* = 0.0544 | B = -2.0172 *p* = 0.4042 | B = 3.5661 *p* = 0.1240 | **B = -13.180 *p* = 0.0215*** |
|  | Head poke duration | | | | | |
|  | Male | | | Female | | |
|  | Rearing  x Brain | Housing  x Brain | R x H x Brain | Rearing  x Brain | Housing  x Brain | R x H x Brain |
| PV count | B = 0.0499 *p* = 0.960 | **B = 2.1413 *p* = 0.0245*** | B = 0.8211 *p* = 0.6905 | B = -1.0063 *p* = 0.340 | B = -1.6558 *p* = 0.0925 | B = -0.6300 *p* = 0.7604 |
| PNN+ PV count | B = -1.7475 *p* = 0.4099 | **B = 3.938 *p* = 0.0475*** | B = -3.007 *p* = 0.5258 | B = -1.456 *p* = 0.5464 | B = -3.1310 *p* = 0.1746 | B = -4.9005 *p* = 0.3160 |
| PNN+ PV intensity | B = 8.9430 *p* = 0.6391 | B = 6.851 *p* = 0.7058 | B = -0.1135 *p* = 0.9978 | B = -11.938 *p* = 0.6262 | B = 20.853 *p* = 0.3945 | B = 96.450 *p* = 0.0506 |
| PV+ PNN intensity | B = -14.019 *p* = 0.6303 | B = 22.807 *p* = 0.4409 | **B = -130.877 *p* = 0.0282*** | B = 22.339 *p* = 0.2530 | B = -11.373 *p* = 0.5568 | B = 44.732 *p* = 0.3469 |

**Supp Table 2.** Two-and three-way interaction regression analyses between measures of infralimbic parvalbumin (PV) cell count, co-labeled perineuronal net (PNN)/PV cell count, PNN+ PV intensity, and PV+ PNN intensity with frequency to open, number of crossings, and head poke duration in the elevated zero maze. Dependent variables for these regressions were Rearing (R), Housing (H), and PNN/PV measure – labeled Brain for simplicity. The unstandardized regression coefficient (B) with the corresponding *p*-value are reported for each regression. *: *p* < 0.05 (also indicated in bold).
